## Supplementary figures and images for "NBR1-mediated selective autophagy of ARF7 modulates root branching"

### Figure EV1

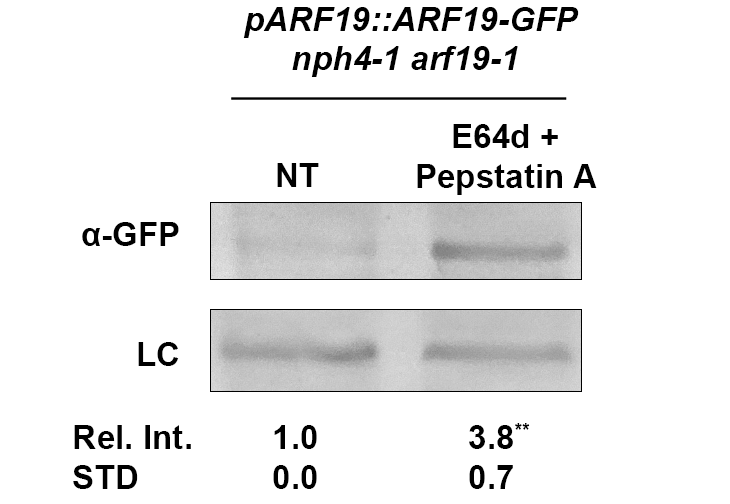

### Figure EV2

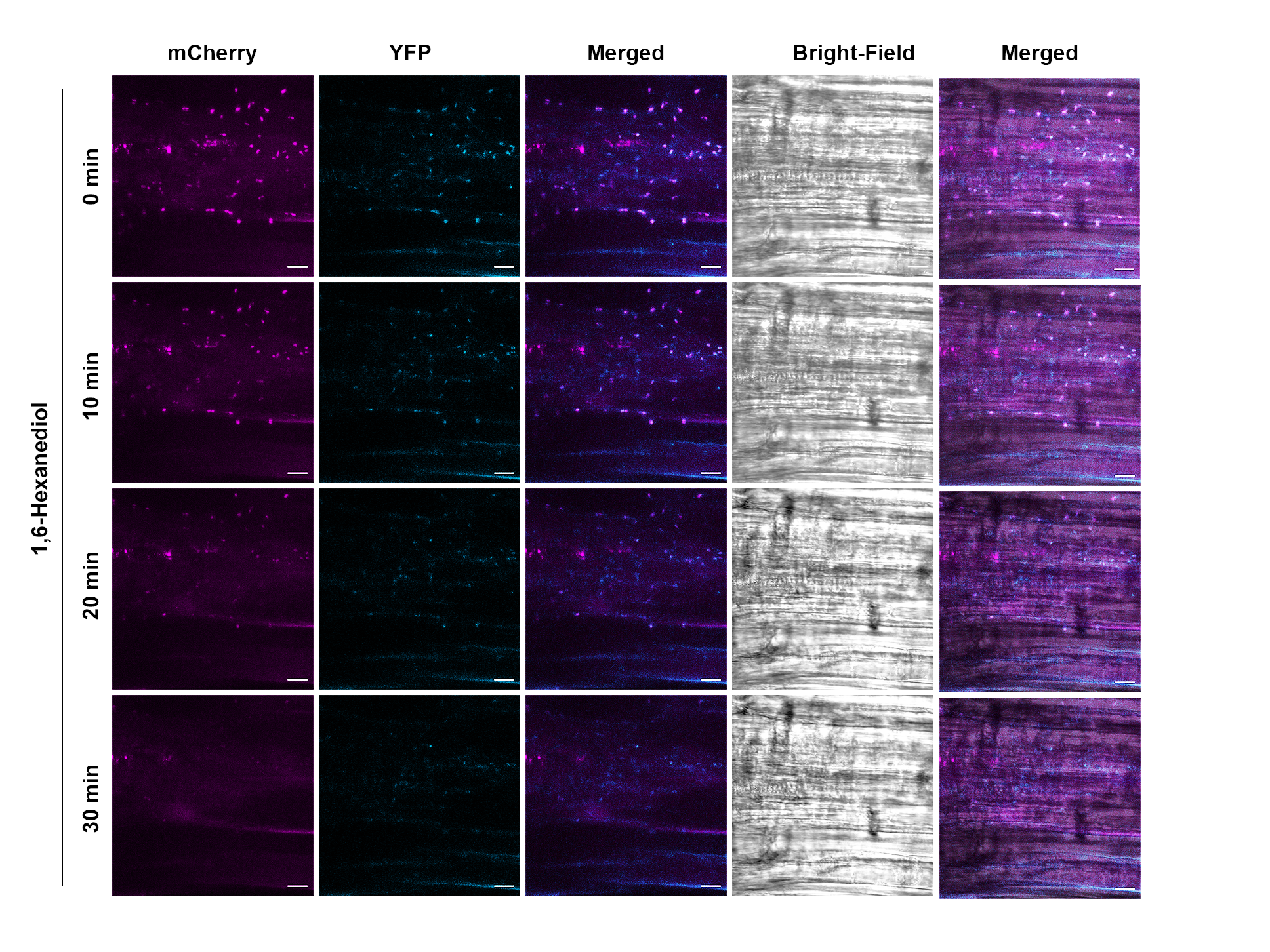

### Figure EV3

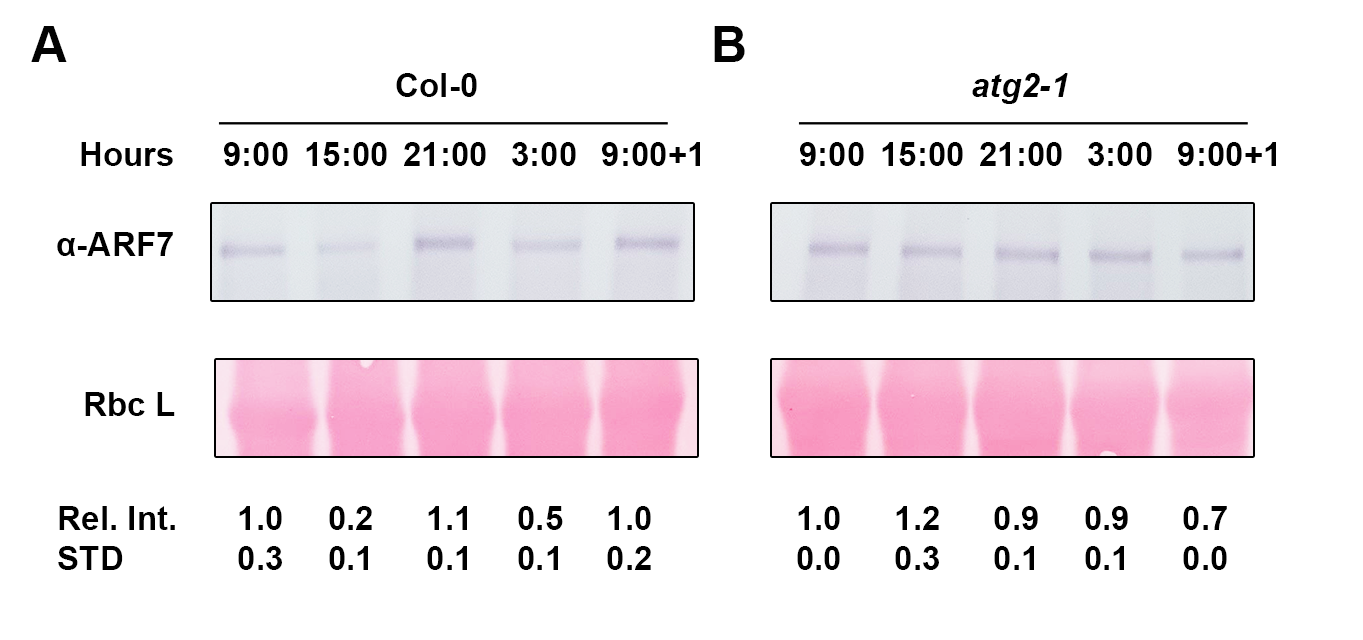

### Figure EV4

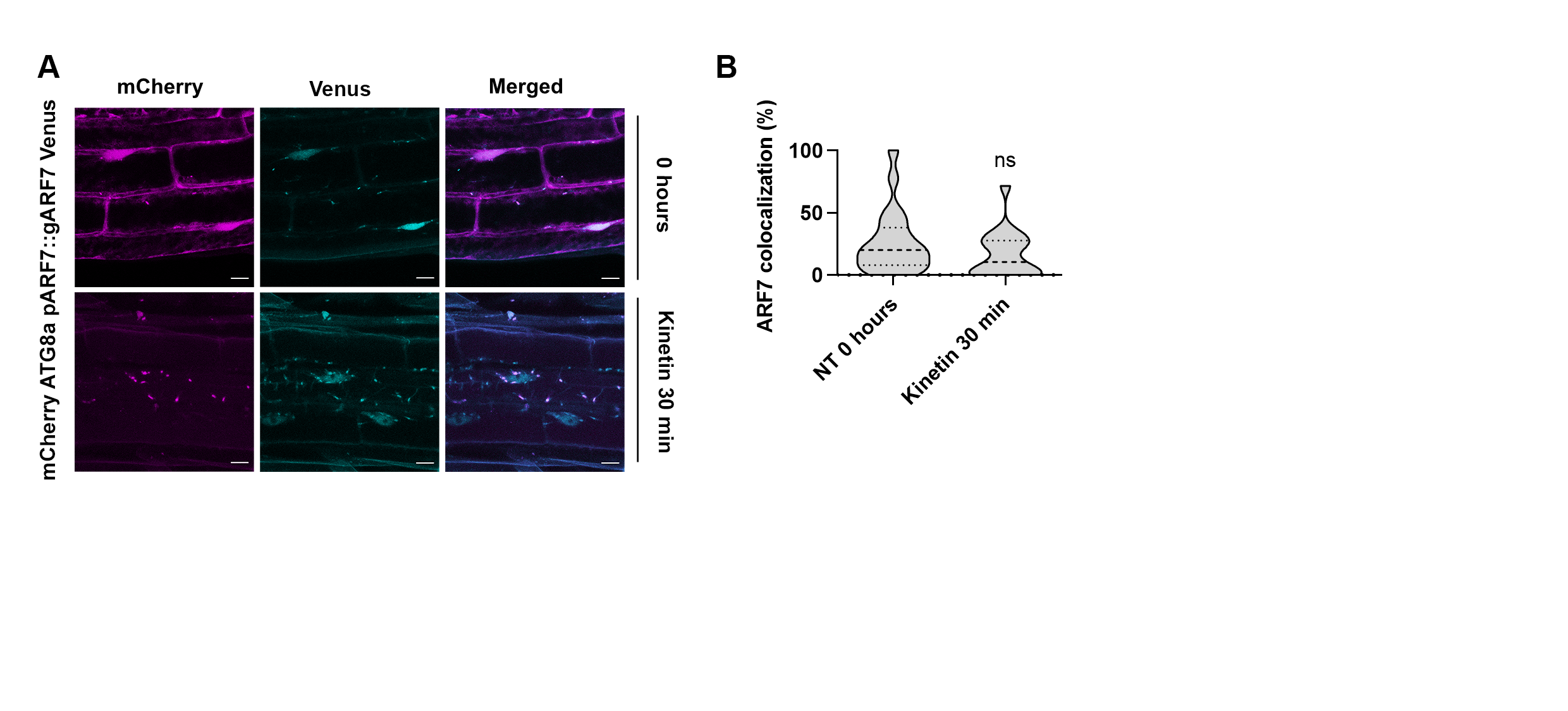
